# Microbial metabolism of complex organic matter across diverse deep-sea ecosystems

**DOI:** 10.1101/2025.11.21.689854

**Authors:** Rikuan Zheng, Chong Wang, Chaomin Sun

**Author notes:** Corresponding authors: Chaomin Sun. E-mail address; Rikuan Zheng. These authors contributed equally to this work (Rikuan Zheng, Chong Wang).

## Abstract

The deep sea is home to a vast and largely unexplored microbial biosphere, along with significant amounts of complex organic matter (COM). However, the ability of deep-sea microbes to metabolize complex organic matter across diverse regions remains poorly understood. Here, we combine 16S rRNA gene amplicon sequencing, metagenomics, and metatranscriptomics to comprehensively characterize prokaryotic communities across different years and habitats (cold seeps, hydrothermal vents, and seamounts). Our results reveal spatio-temporal community heterogeneity driven by geochemical gradients, alongside widespread genomic and transcriptomic potential for COM metabolism. Notably, the PVC (*<u>P</u>lanctomycetota*-*<u>V</u>errucomicrobiota*-*<u>C</u>hlamydiota*) superphylum exhibits extensive polysaccharide degradation capabilities, with *Planctomycetota* strain WC338 and *Lentisphaerota* strain WC36 isolated via laminarin enrichment. Growth and transcriptome data confirm their obligate laminarin dependence and robust degradation capacity, employing distinct enzymes (GH16 and GH2), whose broad distribution across diverse PVC superphylum lineages underscores their prevalence. Furthermore, we demonstrate that laminarin acts as a highly effective selective substrate for enriching and isolating the deep-sea PVC superphylum bacteria. Collectively, these findings delineate specialized adaptations within the PVC superphylum for polysaccharide degradation, significantly expanding our understanding of deep-sea microbial roles in global carbon cycling.

## Introduction

The deep-sea floor encompasses a remarkable range of ecological niches, from the vast, oligotrophic mud plains to highly dynamic, nutrient-rich oases associated with geological features such as cold seeps, hydrothermal vents, and seamounts^1,2^. These benthic oases are characterized by a substantial reservoir of complex organic matter (COM), including proteins, polysaccharides, nucleic acids, and lipids, in addition to humic acid fractions, supporting diverse heterotrophic microbial communities^3–5^. Despite the prevalent energy limitation and characteristically sluggish metabolic rates of the deep subseafloor environment^1,6^, it is a prevailing hypothesis that specific microbial consortia thrive through the efficient degradation of these recalcitrant organic compounds^7^. Indeed, existing evidence indicates that certain isolated deep-sea microbes are equipped with the enzymatic repertoire to catabolize biopolymers^8–10^. However, a systematic, comparative assessment of complex organic matter degradation capacity across diverse deep-sea benthic environments, coupled with targeted pure culture validation, is critically lacking. Bridging this knowledge gap is essential to elucidate the biogeochemical roles and adaptive strategies of these specialized microbial ecosystems.

As one of the most abundant polysaccharides globally, laminarin represents a significant carbon reservoir, comprising approximately 26% of marine particulate organic carbon (POC)^11,12^. Structurally, laminarin is characterized by a β-1,3-D-glucan backbone, often with β-1,6-D-glucan branches, making it a crucial carbon source within marine algae and phytoplankton^13,14^. The rapid degradation of laminarin, particularly during algal blooms, is facilitated by abundant heterotrophic glycoside hydrolases and sugar transporters^15^. Key enzymes involved in laminarin hydrolysis are classified within the Carbohydrate-Active Enzyme (CAZyme) database. Specifically, glycoside hydrolase families GH16, GH17, GH55, GH64, GH81, and GH128 are typically classified as endo-β-1,3-glucanases, whereas GH3 and GH30 possess exo-β-1,3-glucanase activities. Evidence of laminarinase activity is prevalent across various marine strata, from surface waters to deeper water columns and sediments^16,17^, strongly suggesting widespread distribution of laminarin-degrading bacteria and their associated enzymes throughout the world’s oceans. However, despite this ubiquity, a detailed investigation into the capability of deep-sea microorganisms to degrade laminarin remains remarkably scarce. Crucially, there are currently no reports documenting laminarin degradation by purely cultured deep-sea bacteria, particularly those belonging to the PVC superphylum, which constitutes a highly abundant and often dominant phylogenetic lineage in deep-sea environments^18^. Addressing this void requires characterizing the laminarin degradation potential of deep-sea microbes, with a focus on the understudied PVC superphylum, to understand their role in utilizing this widespread carbon source.

The PVC superphylum, encompassing phyla such as *Planctomycetota*, *Verrucomicrobiota*, *Chlamydiota*, *Lentisphaerota*, *Kiritimatiellota*, and several uncultured candidate phyla, represents a remarkable group of bacteria exhibiting traits typically associated with eukaryotes or archaea, or both^19–21^. Among these, *Planctomycetota* have been most extensively isolated and studied. These bacteria are ubiquitous across diverse environments, including freshwater lakes^22^, wetlands^23^, soil^24^, and marine waters and abyssal sediments^25,26^. Notably, *Planctomycetota* are hypothesized to dominate deep-sea sediment communities^27^, with studies in regions like the Gulf of Mexico revealing their substantial relative abundance (up to 28% of bacteria) and apparent involvement in nitrogen cycling and the breakdown of COM delivered as marine snow^18^. However, our understanding of *Planctomycetota*’s functional capabilities in deep-sea environments is severely limited by a profound scarcity of cultured isolates. To date, only two *Planctomycetota* strains have been successfully cultured from deep-sea settings^28–30^. This lack of pure cultures significantly impedes comprehensive characterization of their deep-sea physiology, metabolism, biogeochemical contributions, and ecological roles^31^. Therefore, there is an urgent need to acquire novel deep-sea PVC superphylum isolates, particularly *Planctomycetota*, to enable detailed investigations into their metabolic repertoire, specifically their capacity for degrading complex organic matter such as laminarin.

In this study, our integrated multi-omics and pure culture approach, analyzing samples from diverse deep-sea environments collected in 2018 and 2022, reveals a ubiquitous potential for COM degradation across resident microbial communities. Critically, we experimentally demonstrate a potent laminarin degradation capacity within novel isolates of *Planctomycetota* and *Lentisphaerota*, supported by the widespread conservation and high expression of key laminarinase enzymes. This obligate reliance on laminarin degradation establishes the PVC superphylum as pivotal drivers of carbon cycling within the oligotrophic deep sea. This study thereby unveils a previously unappreciated microbial strategy for nutrient acquisition, highlighting the PVC superphylum’s crucial role in bridging the gap between recalcitrant organic matter and the deep-sea biogeochemical cycle, a key missing piece in our understanding of deep-sea carbon flux.

## Results and discussion

### Microbial community diversity across deep-sea sediment habitats

Deep-sea environment hosts a vast, largely unexplored microbial biosphere, where prokaryotes, endowed with diverse metabolic capabilities, are crucial drivers of organic matter cycling^32,33^. To analyze microbial diversity and composition in different deep-sea sediments, we previously employed 16S rRNA gene amplicon sequencing on sediment samples collected from seven distinct locations and years: 2018 Site F cold seep, 2018 Site H1 hydrothermal vent, 2018 Site H2 hydrothermal vent, 2022 Site F cold seep 1, 2022 Site F cold seep 2, 2022 Haima cold seep, and 2022 Zhenbei seamount (Fig. 1a and Supplementary Data 1)^34^. As shown in Extended Data Figure 1, the microbial communities in the 2018 cold seep and hydrothermal vent samples were predominantly composed of the phyla *Pseudomonadota*, *Chloroflexota*, *Planctomycetota*, *Bacteroidota*, *Acidobacteriota*, and *Bacillota*.

**Figure 1.**
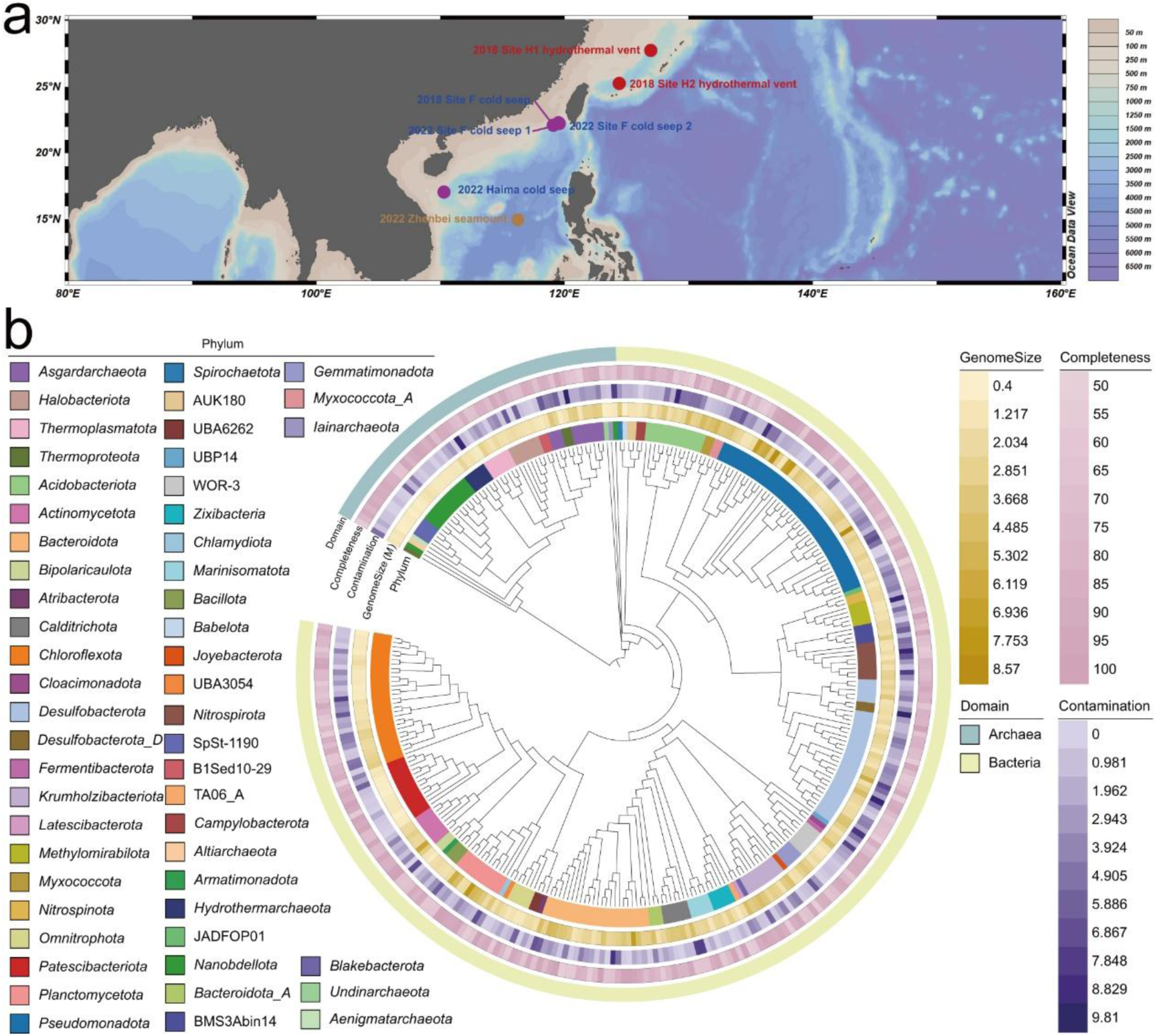
Geographic distribution and phylogenetic analysis of deep-sea metagenome-assembled genomes (MAGs). **a** Geographic distribution of sampling sites across seven deep-sea environments (2018 Site F cold seep, 2018 Site H1 hydrothermal vent, 2018 Site H2 hydrothermal vent, 2022 Site F cold seep 1, 2022 Site F cold seep 2, 2022 Haima cold seep and 2022 Zhenbei seamount). Image generated with Ocean Data View (ODV; Schlitzer, Reiner, https://odv.awi.de, 2021). **b** Phylogenetic tree of MAGs. Innermost ring: phylum-level clades (colored). Second ring: genome size. Third and fourth rings: completeness and contamination levels. Outermost ring: Bacterial (light yellow) and Archaeal (light blue) MAGs.

Notably, *Pseudomonadota* exhibited a pronounced dominance in the cold seep sites, comprising up to 63.1% of the community. In contrast, the two hydrothermal vent sites showed a comparatively lower proportion of *Pseudomonadota* (29.6% on average), with a greater abundance of unclassified bacterial groups (24.1%), alongside *Chloroflexota* (10.7%), and *Planctomycetota* (10.7%). This observed difference in community composition between cold seeps and hydrothermal vents may reflect varying geochemical gradients and nutrient availability^32,35^. As shown in Extended Data Figure 2, the microbial communities in the 2022 cold seep and seamount samples were primarily dominated by the phyla *Chloroflexota*, *Thermodesulfobacteriota*, *Pseudomonadota*, and *Atribacterota*. The abundance of *Thermodesulfobacteriota* (22.5% on average) and *Atribacterota* (19.4% on average) was higher in the cold seep sites compared to the seamount site (0.18% and 0%, respectively). *Thermodesulfobacteriota*, a key group of sulfate-reducing bacteria, are thus expected to thrive in these sulfate-rich environments, which also support the activity of anaerobic methane oxidizers (ANMEs). The microbial communities in these cold seep sediments likely govern the dominant biogeochemical processes through the synergistic microbial sulfate reduction and anaerobic methane oxidation^36^. These findings highlight the substantial heterogeneity in microbial community composition across deep-sea sediment samples, both temporally and spatially.

Metagenomic sequencing of the seven deep-sea sediments yielded 316 high- to medium-quality (completeness ≥50%, contamination ≤10%) microbial metagenome-assembled genomes (MAGs)^37^ after de novo assembly and binning. Clustering at 95% average nucleotide identity (ANI) grouped these into 261 bacterial and 55 archaeal MAGs, representing species-level clusters across 30 bacterial and 10 archaeal phyla (Fig. 1b and Supplementary Data 2). Bacterial MAGs were predominantly derived from dominant lineages, with significant representation from *Pseudomonadota* (n = 43), *Thermodesulfobacteriota* (n = 32), *Chloroflexota* (n = 28), *Bacteroidota* (n = 26), *Acidobacteriota* (n = 13), *Patescibacteriota* (n = 13), *Planctomycetota* (n = 11), and *Nitrospinota* (n = 10). These dominant bacterial phyla prevailed in all seven deep-sea sediments analyzed. Within the Archaea domain, the 55 MAGs were predominantly derived from *Nanobdellota* (n = 14), *Asgardarchaeota* (n = 10), *Halobacteriota* (n = 7), *Thermoplasmatota* (n = 6), *Hydrothermarchaeota* (n = 5), *Thermoproteota* (n = 3), *Aenigmatarchaeota* (n = 1), *Altiarchaeota* (n = 1), *Iainarchaeota* (n = 1), and *Undinarchaeota* (n = 1), along with six unclassified archaeal MAGs. While these archaeal lineages were broadly distributed across the sampled deep-sea sediments, the *Nanobdellota* MAGs exhibited a highly specific distribution, being exclusively recovered from the two hydrothermal vent sediment samples (Supplementary Data 2). *Nanobdellota* was a prominent lineage within the DPANN archaea superphylum, which exclusively the high abundance in hydrothermal vent sediments^4,38^. Together, these findings underscore the pronounced spatial and temporal structuring of deep-sea microbial communities, revealing that distinct assemblages of widespread generalists and niche-specialized taxa drive unique biogeochemical processes in different abyssal habitats.

### Deep-sea microbes possess substantial potential for COM metabolism

To estimate the genomic potential for degrading COM, we screened these MAGs for genes encoding enzymes involved in the breakdown of proteins, polysaccharides, nucleic acids, and lipids. Initially, the majority of MAGs encoded extracellular peptidases, predominantly found in dominant phyla such as *Pseudomonadota*, *Thermodesulfobacteriota*, *Bacteroidota*, *Halobacteriota*, and *Methylomirabilota* (Fig. 2 and Supplementary Data 3). Analysis of their protein transport systems indicated that most MAGs possess transporters for polar amino acids, branched-chain amino acids, peptides, and oligopeptides. The presence of genes encoding amino acid degradation enzymes (*gdh*, *ior*, *kor*, *por*, and *vor*) suggests that these MAGs can likely utilize environmental proteins as energy sources. Remarkably, a significant proportion of our recovered MAGs, particularly from these cold seep, hydrothermal vent, and seamount sediments, encode most amino acid transport and degradation pathways, contrasting sharply with previous reports of widespread absence of such pathways in MAGs from gas hydrate-bearing subseafloor and the Helgoland mud are^39,40^.

**Figure 2.**
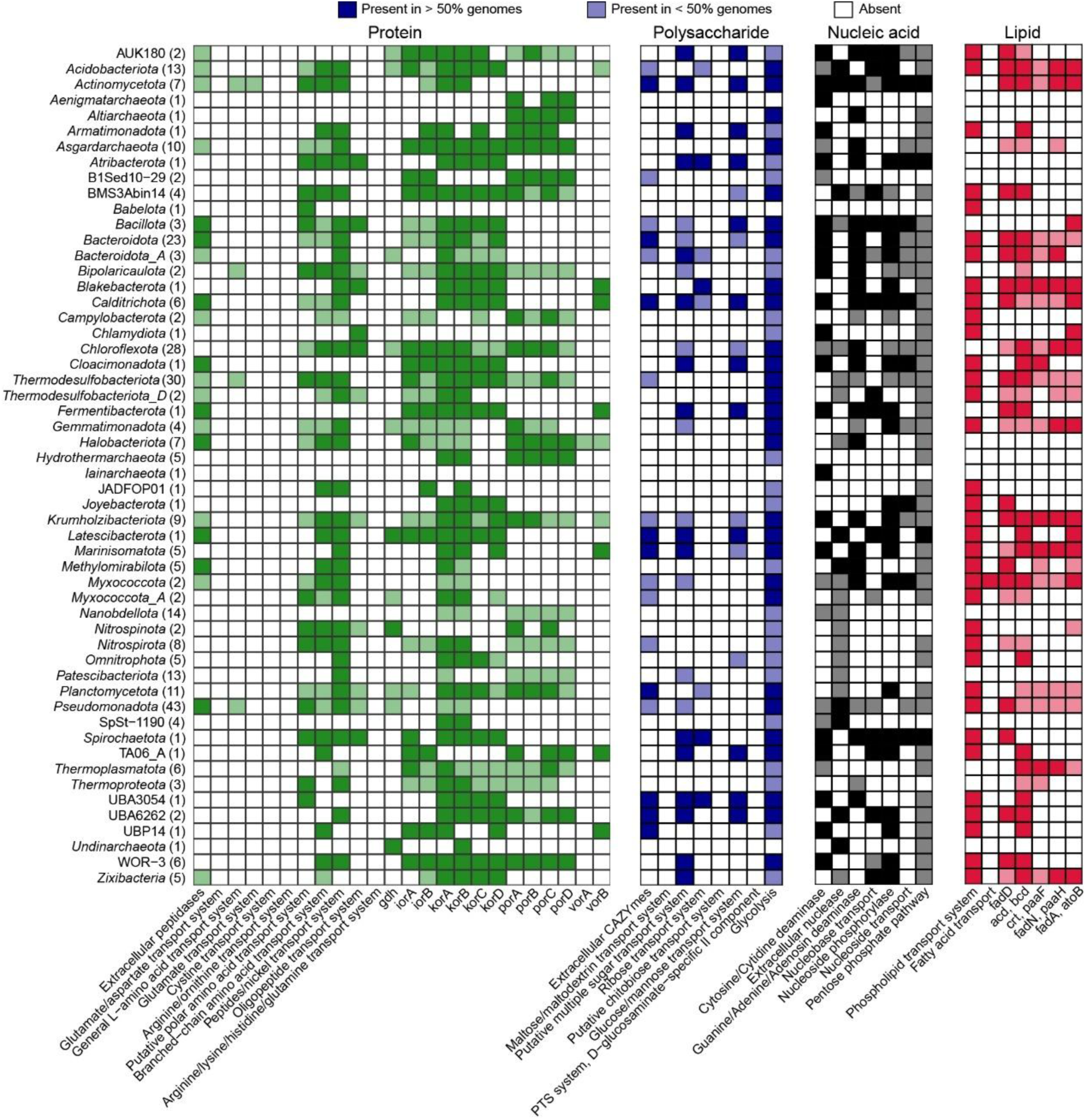
Distribution of key catabolic genes across deep-sea MAGs reveals diverse degradation capabilities of complex organic matter. Key genes responsible for the catabolism of proteins, polysaccharides, nucleic acids, and lipids were detected in MAGs. Color intensity indicates gene prevalence within each cluster: dark shading denotes presence in >50% of MAGs, and light shading indicates presence in <50% of MAGs. The total number of MAGs per cluster is shown in parentheses. Pathway identification was performed with METABOLIC.

Subsequently, regarding polysaccharide degradation, we detected 17,093 putative CAZymes within 316 MAGs (Fig. 2 and Supplementary Data 4). Among these, glycoside hydrolases (GHs) were the most abundant (45.7%), followed by glycosyltransferases (GTs, 39.2%), carbohydrate-binding modules (CBMs, 9.5%), and carbohydrate esterases (CEs, 4.3%). Crucially, most MAGs encoded diverse extracellular CAZymes capable of cleaving a wide range of carbohydrates, coupled with numerous sugar transport systems for the uptake of oligo- and monomers. These breakdown products can subsequently enter the ubiquitous glycolysis pathway. However, in stark contrast to the prevalent polysaccharide degradation capabilities observed in some bacterial lineages, the vast majority of archaeal MAGs in our dataset consistently lacked genes encoding extracellular CAZymes (Supplementary Data 4). This genomic deficit suggests a limited capacity for the direct extracellular hydrolysis and utilization of complex polysaccharides by deep-sea archaea. Instead, our analysis revealed that nearly all archaeal and bacterial MAGs harbor genes involved in glycolysis, indicating a strong potential for fermentative metabolism of simpler sugars such as glucose, which was consistent with previous studies^39^.

While nucleic acids are recognized as a fundamental trophic resource and energy source in the deep sea^41^, their degradation potential across the microbial biosphere has remained largely an enigma. Prior investigations have primarily focused on specific bacterial isolates, such as our previous demonstration of extracellular DNA (eDNA) utilization by deep-sea *Xianfuyuplasma coldseepsis* zrk13^10^. Our results showed that the genomic potential for nucleic acid degradation was distributed across both bacterial and archaeal lineages (Fig. 2 and Supplementary Data 5). These MAGs encoded a suite of genes facilitating efficient heterotrophic nutrition. Key capabilities included extracellular nucleases and phosphohydrolases for liberating nucleosides and nucleobases, complemented by transporters for their uptake and dedicated pathways for purine and pyrimidine degradation. The resultant ribose could be channeled into the pentose phosphate pathway. Furthermore, a significant prevalence of genes for lipid catabolism was observed, encompassing phospholipid and fatty acid transporters along with complete beta-oxidation machineries for efficient energy generation (Fig. 2 and Supplementary Data 6). However, certain archaeal lineages, such as *Nanobdellota*, *Aenigmatarchaeota*, *Altiarchaeota*, *Halobacteriota*, *Hydrothermarchaeota*, and *Iainarchaeota*, lacked the genetic machinery for fatty acid degradation. Collectively, these findings indicate that the majority of deep-sea microorganisms possess a substantial capacity to degrade COM.

### Ubiquitous polysaccharide degradation potential in deep-sea microbes

In marine ecosystems, polysaccharides, primarily derived from the cell wall constituents of phytoplankton and macroalgae, represent a substantial carbon input into the deep sea upon sedimentation^9,42^. These recalcitrant molecules are considered a major carbon source for benthic microbial communities. However, current understanding of polysaccharide degradation potential within deep-sea environments is notably limited, largely stemming from the prevailing paradigm that primarily associates specialized polysaccharide-degrading capabilities with specific bacterial lineages, such as the *Bacteroidota* phylum^8,9^. This dominant focus has arguably overlooked the broader metabolic repertoire of other deep-sea microorganisms. To elucidate the metabolic potential of deep-sea microorganisms for polysaccharide degradation and utilization, metagenomic and metatranscriptomic analyses were performed on four deep-sea sediment samples collected in 2022, focusing on CAZyme repertoires. Our screen for CAZymes revealed their widespread distribution across the generated MAGs, encompassing major families including GHs, GTs, PLs, CEs, AAs, and CBMs (Fig. 3 and Supplementary Data 7). To ascertain the functional relevance of these genes in the deep-sea *in situ* environments, metatranscriptomic data demonstrated high expression levels of many CAZyme families in most MAGs, particularly for GHs and GTs, suggesting active polysaccharide degradation pathways (Supplementary Datas 8-13). These actively expressed CAZymes were predominantly associated with dominant bacterial lineages such as *Thermodesulfobacteriota*, *Bacteroidota*, *Halobacteriota*, *Planctomycetota*, and *Omnitrophota* (Fig. 3). Notably, *Planctomycetota* and *Omnitrophota* belong to the PVC superphylum, a group previously reported for its potential to degrade polysaccharides in other environments^43,44^. Our findings demonstrate active polysaccharide degradation in deep-sea PVC superphylum bacteria, revealing their significant metabolic contribution. This discovery extends our understanding of the ecological roles of this understudied the PVC superphylum in marine carbon cycling and highlights the diverse enzymatic repertoire of deep-sea microbial communities for breaking down complex biopolymers.

**Figure 3.**
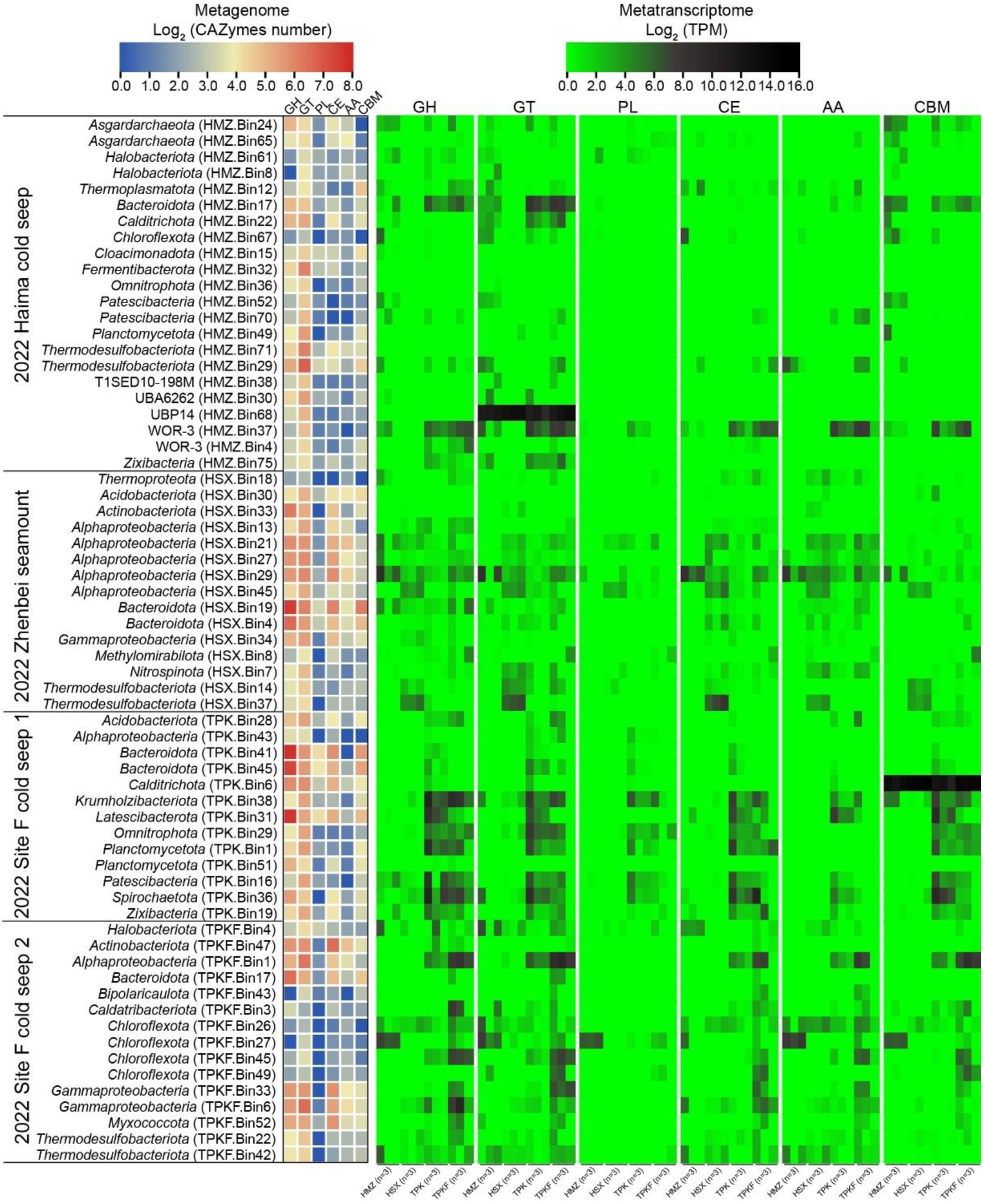
Abundance and expression of carbohydrate-active enzymes across deep-sea microbial communities. The left panel shows the abundance of key carbohydrate-active enzyme (CAZyme) families (GH, glycoside hydrolase; GT, glycosyltransferase; PL, polysaccharide lyase; CE, carbohydrate esterase; AA, auxiliary activity; CBM, carbohydrate-binding module) within deep-sea MAGs. MAGs are organized into distinct phylogenetic clusters from four deep-sea sites: 2022 Haima cold seep, 2022 Zhenbei seamount, 2022 Site F cold seep 1, and 2022 Site F cold seep 2. The right panel displays the expression levels of CAZyme-encoding genes within MAGs, quantified as log_2_-transformed transcripts per million (log_2_TPM) from metatranscriptomic data.

### Obligate laminarin utilization by deep-sea PVC superphylum bacteria

Based on their polysaccharide degradation potential, we developed an optimized enrichment strategy using laminarin (Fig. 4a). This selective cultivation yielded PVC superphylum bacteria, including *Planctomycetota* and *Lentisphaerota* representatives, from which high-growth strains WC338 and WC36 were isolated for further study. Preliminary phenotypic characterization via transmission electron microscopy (TEM) revealed that strain WC338 possesses a single polar flagellum and a spherical morphology (average diameter of 1.0 µm) (Fig. 4b), whereas strain WC36 presented an oval shape (approximately 1.0 µm in size) and appeared to lack flagellar structures (Fig. 4c). Phylogenetic analyses based on 16S rRNA gene sequences and sequence identity calculations using NCBI BLAST revealed the distinct taxonomic placements of deep-sea isolates WC338 and WC36. Strain WC338 exhibited high sequence identity to members of the genus *Poriferisphaera*, including *P. corsica* KS4 (98.06%) and *P. heterotrophicis* ZRK32 (97.19%), but fell below the species demarcation threshold (98.65%), thus strongly suggesting it represents a novel species within *Poriferisphaera*. In parallel, strain WC36 displayed a unique phylogenetic position within the class *Lentisphaeria*, forming a deeply branching lineage equidistant from the orders *Lentisphaerales* and *Victivallales*. This significant phylogenetic divergence, coupled with sequence divergence thresholds for order delineation, indicates that strain WC36 may represent a novel order within *Lentisphaeria*, thereby expanding the known taxonomic diversity of this understudied class. In parallel, strain WC36 showed high sequence identity to members of the order *Lentisphaerales* (e.g., *Victivallis lenta* BBE-744-WT-12, 85.95%; *V. vadensis* Cello, 85.14%) and the order *Victivallales* (e.g., *Lentisphaera profundi* SAORIC-696, 82.95%; *L. araneosa* HTCC2155, 82.76%). Phylogenetically, strain WC36 formed a distinct, deeply branching lineage within the class *Lentisphaeria*, equidistant from these two orders. Given the typical order delineation threshold (around 83.55%) and its significant phylogenetic distance, strain WC36 likely represents a novel order, thereby expanding the taxonomic diversity of this understudied class.

**Figure 4.**
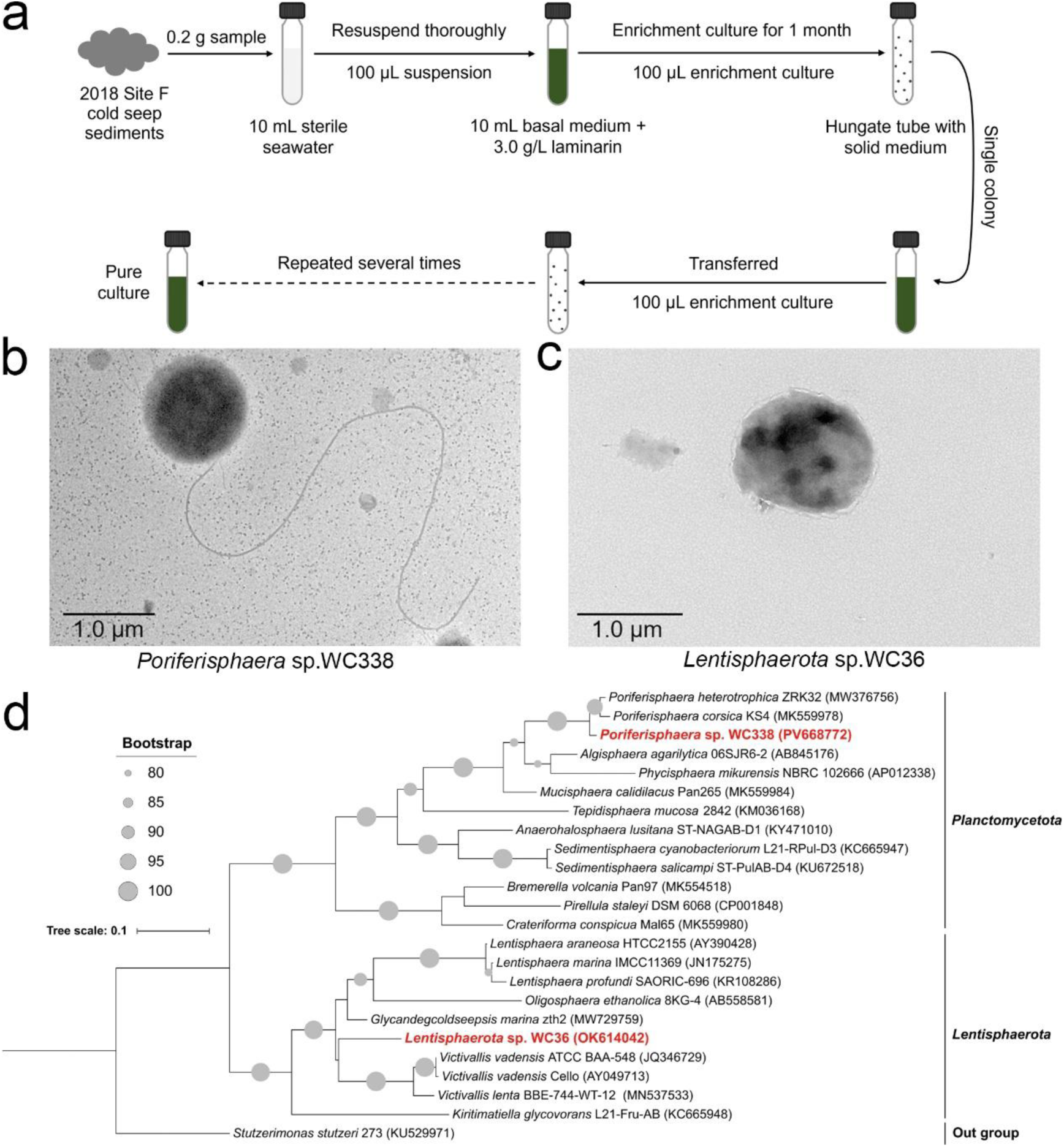
Isolation, morphology, and phylogenetic analysis of cultured deep-sea PVC superphylum bacteria. **a** Isolation strategy for PVC superphylum bacteria. **b, c** Transmission electron microscopy (TEM) micrographs of *Poriferisphaera* sp. WC338 (C) and *Lentisphaerota* sp. WC36 (D). **d** Phylogenetic tree of *Poriferisphaera* sp. WC338 and *Lentisphaerota* sp. WC36. Strain WC338 is placed within the phylum *Planctomycetota* and strain WC36 within the phylum *Lentisphaerota*, based on 16S rRNA gene sequences. Bootstrap support values (>80%) are indicated at the base of each node by gray dots. Scale bar represents 0.1 substitutions per nucleotide position.

NCBI BLASTn analysis revealed that strain WC338 exhibited high sequence identity to *Poriferisphaera corsica* KS4 (98.06%), *Poriferisphaera heterotrophicis* ZRK32 (97.19%), *Mucisphaera calidilacus* Pan265 (98.09%), *Algisphaera agarilytica* 06SJR6-2 (88.73%), and *Phycisphaera mikurensis* NBRC 102666 (85.55%). For strain WC36, the closest relatives identified were *Victivallis lenta* BBE-744-WT-12 (85.95%), *Victivallis vadensis* Cello (85.14%), *Lentisphaera profundi* SAORIC-696 (82.95%), and *Lentisphaera araneosa* HTCC2155 (82.76%). To further confirm the taxonomic affiliations of strains WC338 and WC36, phylogenetic trees were constructed using 16S rRNA gene sequences (Fig. 4d). Robustly supported by the maximum likelihood phylogenetic tree, strain WC338 was classified as a member of *Poriferisphaera*. Within this genus, it formed a distinct monophyletic clade with *P. corsica* KS4 and *P. heterotrophicis* ZRK32, positioned as an independent lineage within the broader *Poriferisphaera* radiation. Crucially, the sequence identity of WC338 to its closest *Poriferisphaera* relatives (e.g., *P. corsica* KS4 at 98.06%) falls below the commonly accepted threshold for species demarcation (98.65%)^45^. T Phylogenetic analysis and sequence divergence provide robust evidence that strain WC338 constitutes a novel species of *Poriferisphaera*. In parallel, the phylogenetic analysis of strain WC36 revealed its placement within the class *Lentisphaeria*. Strain WC36 formed a distinct, deeply branching lineage that occupied a position equidistant from both the order *Lentisphaerales* and the order *Victivallales*. Given the taxonomic threshold for order delineation (typically around 83.55%)^46^, the significant phylogenetic distance and distinct branching pattern observed for strain WC36 suggest that it may represent a novel order within the class *Lentisphaeria*. This finding significantly expands the taxonomic diversity of this understudied class.

Metabolic analysis confirmed the direct utilization of laminarin by these two PVC superphylum strains. When cultured in basal medium where laminarin was supplied as the sole carbon source, both strains reached stationary phase within 5 to 6 days (OD_600_∼0.16) (Fig. 5a-b). Crucially, this growth correlated directly with laminarin consumption, evidenced by a significant decrease in laminarin relative content from 100% down to 55.7% (strain WCC338) and 66.5% (strain WC36). These results robustly demonstrate that these PVC superphylum bacteria can effectively catabolize laminarin to sustain growth and maintain viability. Furthermore, the growth of both strains exhibited strict nutritional dependence on laminarin. Supplementation of the basal medium with common energy sources, including glucose, N-acetylglucosamine, yeast extract, or peptone, failed to support measurable growth (Fig. 5c-d). Intriguingly, while N-acetylglucosamine is a frequently utilized substrate by *Planctomycetota* bacteria^28^, strain WC338 could not utilize it. Laminarin, a key component of the marine carbon cycle, constitutes approximately 26% of the marine particulate organic carbon pool, underscoring its substantial contribution^11,12,14^. Therefore, this strict dependence on laminarin coupled with the inability to assimilate simpler carbon sources like N-acetylglucosamine, strongly suggests that these deep-sea PVC superphylum bacteria possess unique, highly specialized metabolic pathways adapted specifically for laminarin assimilation, potentially reflecting their specialized niche in the deep-sea carbon cycle.

**Figure 5.**
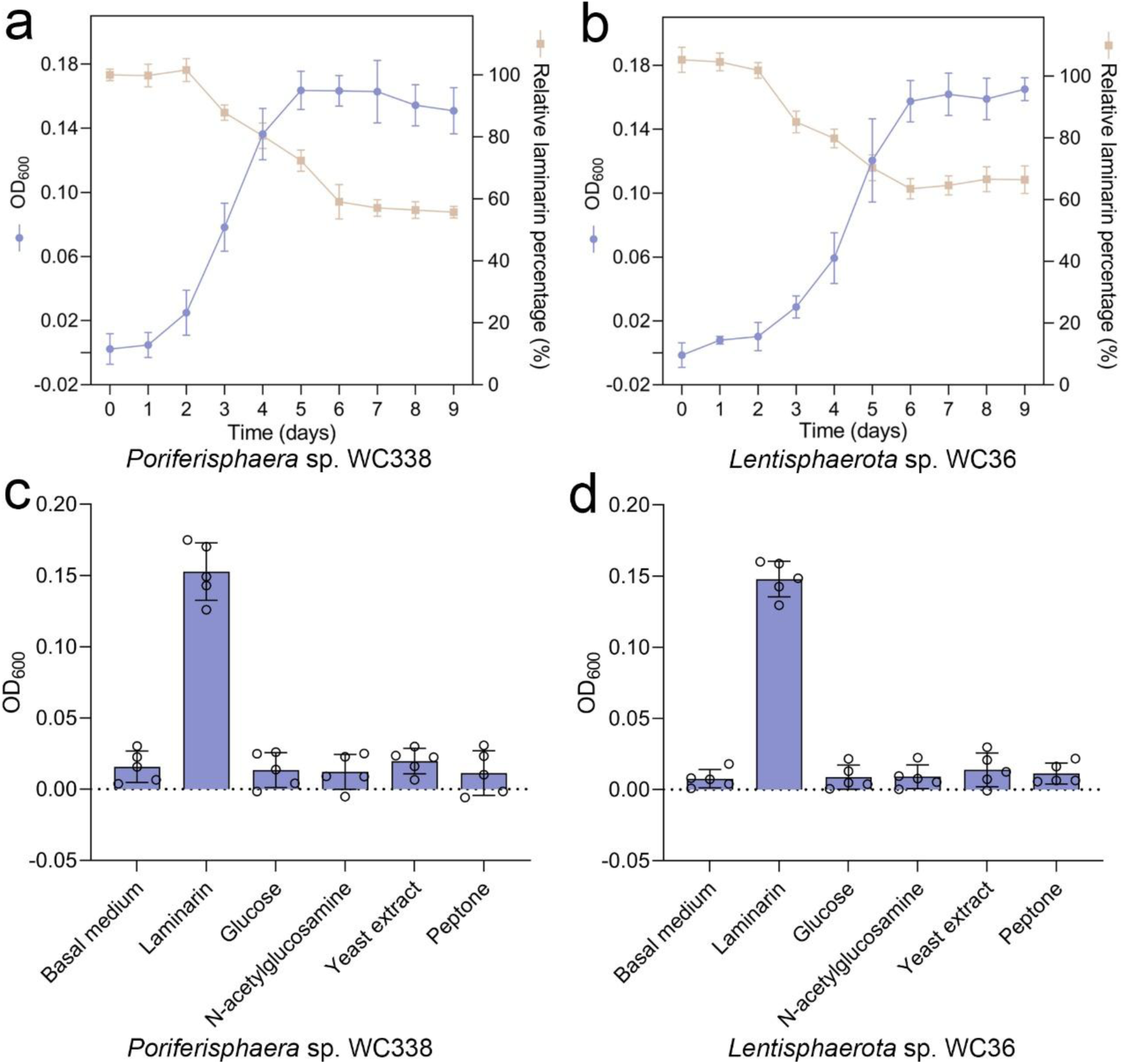
Obligate laminarin dependency drives growth of deep-sea PVC superphylum strains WC338 and WC36. **a** Growth of strain WC338 is coupled with laminarin consumption. Growth curves (Purple line, OD_600_) and relative laminarin concentrations (Orange-yellow line, relative percentage) of strain WC338. **b** Growth of strain WC36 is coupled with laminarin consumption. Growth curves (Purple line, OD_600_) and relative laminarin concentrations (Orange-yellow line, relative percentage) of strain WC36. **c, d** Growth of strains WC338 (C) and WC36 (D) on various carbon sources. Bacterial growth was assessed in basal medium alone or supplemented with 3.0 g/L of either laminarin, glucose, N-acetylglucosamine, yeast extract, or peptone as the sole carbon source.

### The strategy for laminarin catabolism in deep-sea PVC superphylum bacteria

To decipher the mechanisms underlying laminarin degradation and utilization by strains WC338 and WC36, we first performed a systematic analysis of their CAZymes. Strain WC338 exhibited a substantially larger complement of glycoside hydrolases (GHs) and glycosyltransferases (GTs) compared to strain WC36 (Figs. 6a-b). Notably, WC338 possessed canonical laminarin-degrading enzymes, specifically GH16 and GH13^11^, which were absent in strain WC36. To further probe the metabolic characteristics of strains WC338 and WC36, we performed transcriptomic analyses under laminarin induction. A volcano plot analysis revealed significant transcriptional upregulation of 255 genes in strain WC338 (Fig. 6c), with our analysis focusing on genes encoding GHs, sugar transporters, and components involved in flagellar motility. Consistent with its genomic potential and laminarin degradation phenotype, strain WC338 showed significant upregulation of genes encoding GH16 and GH13 (Fig. 6d). Additionally, genes encoding other GH families (GH1, GH2, GH4, GH20) and numerous sugar transporter proteins (Fig. 6e), crucial for importing breakdown products, were also upregulated. Furthermore, a substantial cluster of genes related to flagellar motility and chemotaxis exhibited elevated expression (Fig. 6f; given that marine bacteria are known to exhibit chemotaxis towards laminarin^47^, this upregulation potentially facilitates rapid capture of laminarin, a role further suggested by the single polar flagellum in WC338. In parallel, under laminarin induction, strain WC36 displayed significant upregulation of 97 genes (Fig. 6g). Intriguingly, although lacking canonical laminarin-degrading enzymes like GH16 and GH13 found in WC338, strain WC36 showed significant upregulation of genes encoding GH2 (Fig. 6h). GH2 enzymes are known to hydrolyze *β*-1,4-glycosidic bonds^48^, a prevalent linkage in laminarin^49^, suggesting that WC36 may employ a distinct GH2-based pathway for laminarin breakdown, a mechanism not previously reported for this class of enzymes in laminarin degradation. Consistent with this, genes encoding sugar transporters were also significantly upregulated in WC36 (Fig. 6i), indicating efficient uptake of breakdown products. Interestingly, while strain WC36 appears to lack flagellar structures, genes associated with flagellar protein synthesis were significantly upregulated (Fig. 6j), suggesting that these genes might potentially regulate other cellular functions or adaptation mechanisms. Collectively, these findings reveal differential strategies employed by closely related deep-sea PVC superphylum bacteria for laminarin catabolism, highlighting the metabolic plasticity and specialized adaptations of these lineages in the deep-sea environment.

**Figure 6.**
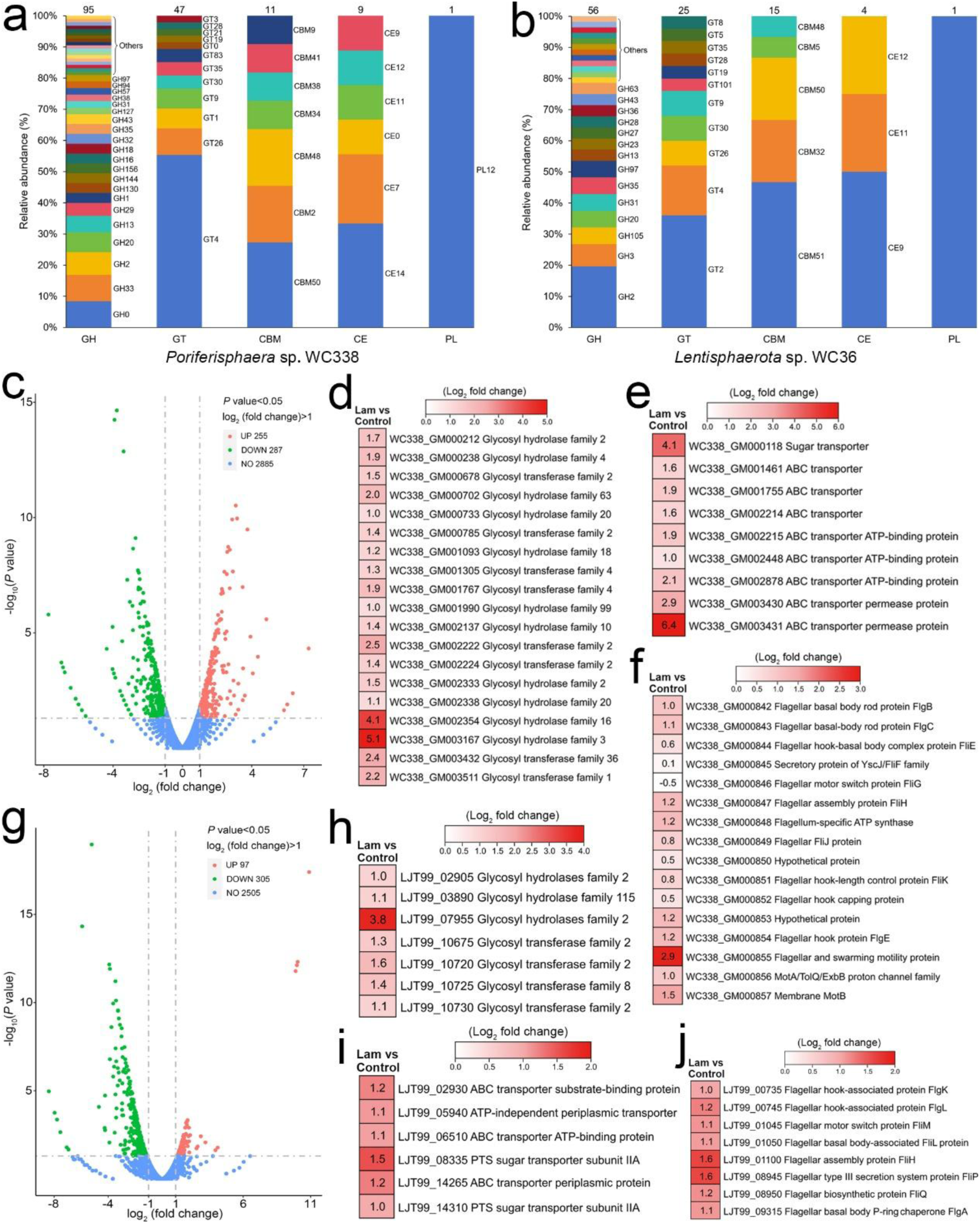
CAZymes composition and transcriptomic analyses of *Poriferisphaera* sp. WC338 and *Lentisphaerota* sp. WC36. Statistical analysis of the types and numbers of CAZymes in *Poriferisphaera* sp. WC338 (**a**) and *Lentisphaerota* sp. WC36 (**b**). **c** Volcano plot showing the number of significantly differentially expressed genes in strain WC338 cultured with 3.0 g/L laminarin. Transcriptomic analysis of strain WC338 cultured with 3.0 g/L laminarin reveals the expression of genes encoding GHs (**d**), sugar transporters (**e**), and components involved in flagellar motility (**f**). **g** Volcano plot showing the number of significantly differentially expressed genes in strain WC36 cultured with 3.0 g/L laminarin. Transcriptomic analysis of strain WC36 cultured with 3.0 g/L laminarin reveals the expression of genes encoding GHs (**h**), sugar transporters (**i**), and components involved in flagellar motility (**j**). “Lam” indicates strains WC338 and WC36 grown in the basal medium with 3.0 g/L laminarin; “Control” indicates strains WC338 and WC36 grown in the basal medium with 0.05 g/L laminarin. These numbers in panels D-F and H-J represent the fold change of gene expression (by using the log_2_ value).

### Ubiquitous laminarin catabolism by PVC superphylum bacteria across diverse environments

Extending the investigation to the broader PVC superphylum reveals widespread laminarin-degrading potential. Building upon the discovery that strains WC338 and WC36 exhibit efficient laminarin degradation, we hypothesized that this capability might be a common trait across the PVC superphylum. To test this, we conducted a comprehensive genomic survey of laminarin-relevant glycoside hydrolase (GH) genes, specifically focusing on GH16 and GH2 due to their direct role in hydrolyzing the β-1,4-glycosidic bonds characteristic of laminarin, as well as other GH families (GH3, GH17, GH55, GH64) found in related taxa. Our analysis encompassed 171 pure cultures of bacteria belonging to the PVC superphylum, including members from the phyla *Planctomycetota*, *Verrucomicrobia*, *Lentisphaerota*, and *Kiritimatiellota*. These isolates were derived from diverse environmental niches worldwide, including marine sediments, water columns, terrestrial soils, and algal samples (Supplementary Data 14), underscoring the broad ecological distribution of this bacterial group (Fig. 7a). Our genomic analysis revealed that genes encoding GH16 and GH2 are widely distributed within the PVC superphylum bacteria (Fig. 7b and Supplementary Data 15), suggesting a prevalent capacity for laminarin utilization. In contrast, genes for GH3, GH17, GH55, and GH64 were found in a smaller subset of these bacteria, indicating more specialized or context-dependent roles. The high prevalence of GH16 and GH2, particularly in isolates from diverse marine environments, strongly supports the hypothesis that laminarin serves as a significant and widespread carbon source in various deep-sea and marine ecosystems^14^, and that PVC superphylum bacteria have evolved robust enzymatic machinery to exploit it. This broad distribution of laminarin-degrading genes highlights the crucial role of the PVC superphylum bacteria in the marine carbon cycle and underscores their potential importance in the breakdown of algal biomass, a major component of oceanic organic matter ^13^.

**Figure 7.**
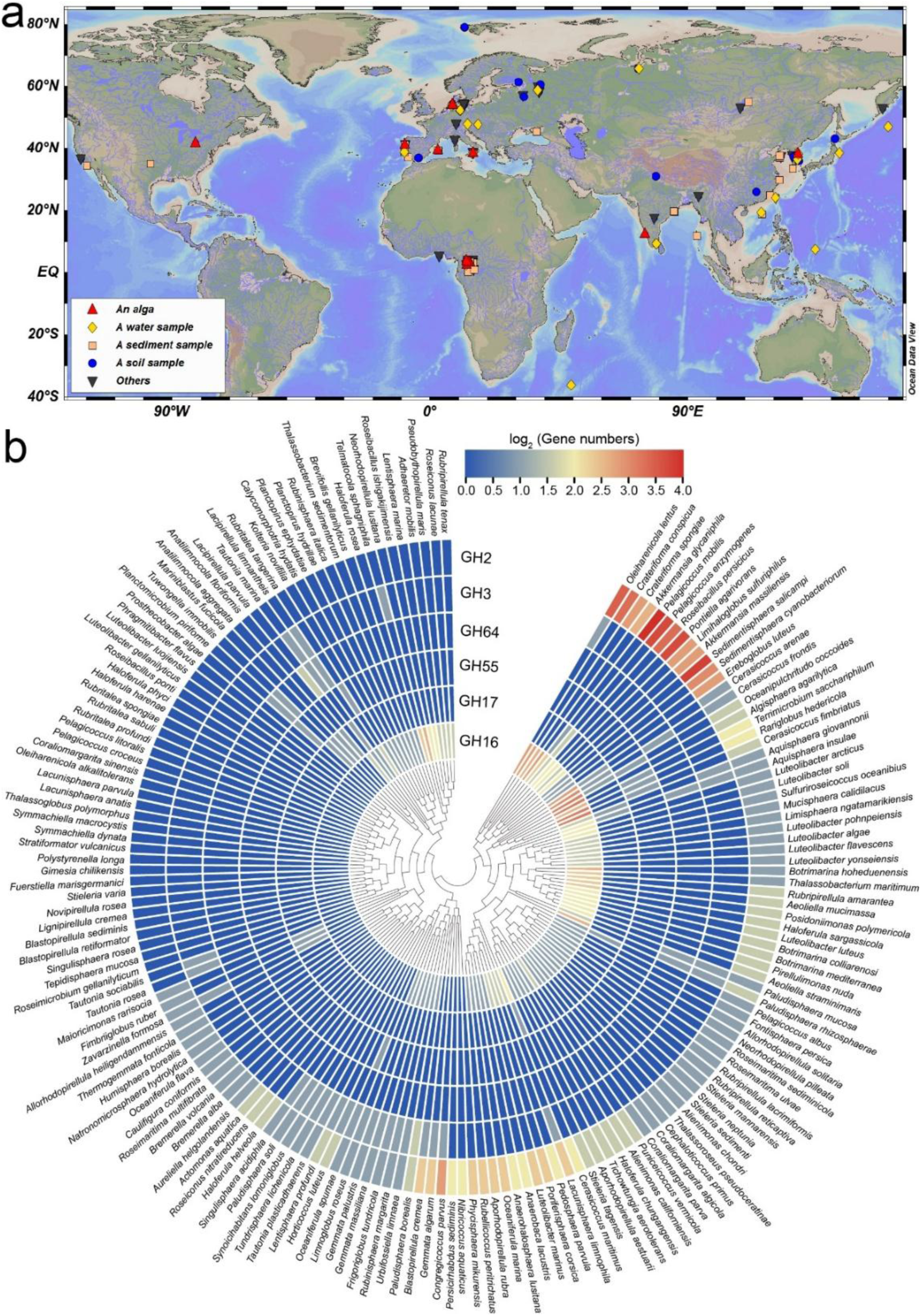
Geographic distribution and glycoside hydrolase gene profiles of PVC superphylum bacteria from diverse environmental sources. **a** Geographic distribution of PVC superphylum bacteria isolates from various environmental sources. Isolates are classified by environmental source: alga (red triangle), water (yellow diamond), sediment (orange square), soil (blue circle), and others (gray inverted triangle). **b** Clustered heatmap of laminarin-degrading GHs in the PVC superphylum bacteria. A circular heatmap displays the log_2_-transformed GH gene copy numbers associated with laminarin degradation (GH16, GH3, GH17, GH55, GH64, and GH2) for 171 PVC bacterial genomes, arranged according to hierarchical clustering based on GH gene content.

### Laminarin facilitates the isolation of deep-sea PVC superphylum bacteria

The ability of laminarin to selectively enrich *Planctomycetota* strain WC338 and *Lentisphaerota* strain WC36 from deep-sea cold seep sediments suggests that laminarin-based cultivation strategies might be broadly applicable to the isolation of other recalcitrant bacterial taxa across diverse deep-sea environments. To test this hypothesis, we performed enrichment cultures of sediment samples from hydrothermal vents and seamounts using basal medium with laminarin supplementation, for 30 days. Enrichment cultures of basal medium supplemented with laminarin led to the isolation of novel pure strains, including one *Planctomycetota* (strain WC47) from hydrothermal vent sediment and one *Lentisphaerota* (strain WC29) from seamount sediment (Extended Data Figure 3), indicating that laminarin is a very effective material for enriching and isolating deep-sea PVC superphylum bacteria. To date, only two *Planctomycetota* strains, (*Gimesia benthica* E7 and *Poriferisphaera heterotrophicis* ZRK32)^29,30^ and one *Lentisphaerota* strain (*Lentisphaera profundi* SAORIC-696)^50^ have been successfully isolated and characterized from deep-sea habitats, highlighting the challenges associated with accessing and studying these deep-sea PVC superphylum bacteria. The successful isolation of diverse PVC superphylum members via laminarin enrichment supports previous metagenomic and metatranscriptomic evidence suggesting their metabolic capability to degrade laminarin.

## Methods

### Sample collection

2018 Site F cold seep, 2018 Site H1 and H2 hydrothermal vent samples were collected by *RV KEXUE* in 2018 from two key regions: the South China Sea and the Okinawa Trough (Western Pacific). Deep-sea sediment samples (2022 Site F cold seep 1, 2022 Site F cold seep 2, 2022 Haima cold seep, and 2022 Zhenbei seamount) were collected from South China Sea during July 2022 using the *RV KEXUE*. These sediment samples, described in Supplementary Data 1, were collected from geographically distinct locations: 2018 Site F cold seep (119°10′14.64″ E, 22°03′57.10″ N, depth 1,146 m), 2018 Site H1 hydrothermal vent (126°53′50.24″ E, 27°47′11.11″ N, depth 2,468 m), 2018 Site H2 hydrothermal vent (124°22′24.85″ E, 25°15′47.45″ N, depth 2,194 m), 2022 Site F cold seep 1 (119°10′14.64″ E, 22°03′57.10″ N, depth 1,140 m), a2022 Site F cold seep 2 (sediment site, 119°34′48.42″ E, 22°14′35.65″ N, depth 1,140 m), 2022 Haima cold seep (110°21′00.80″ E, 17°04′17.49″ N, depth 1,356 m), and 2022 Zhenbei seamount (116°15′36.44″ E, 15°04′16.08″ N, depth 2,469 m). Sediment samples were immediately stored at −80 °C following rapid collection.

### Amplicon and metagenomic sequencing

To understand the microbial community structure of these sediment samples, 16S rRNA gene sequencing analysis was performed. For 2018 Site F cold seep, 2018 Site H1 hydrothermal vent, and 2018 Site H2 hydrothermal vent samples, these sediment samples were dispatched to Novogene (Tianjin, China) for operational taxonomic unit (OUT) sequencing, as previously described^34^. For 2022 Site F cold seep 1, 2022 Site F cold seep 2, 2022 Haima cold seep, and 2022 Zhenbei seamount samples, the total DNA was extracted using a E.Z.N.A.® soil DNA Kit (Omega Bio-tek, USA), as previously described^51^. For metagenomic sequencing analysis, total DNA was extracted from different sediment samples (20 g each) using the E.Z.N.A.® Stool DNA Kit (Omega Bio-tek, USA). The comprehensive sequencing and analysis protocols were previously established^52^.

### Taxonomic assignments of MAGs

Genome taxonomy of MAGs was assigned using GTDB-Tk v2.1.0^53^. The concatenated alignments of 120 bacterial or 122 archaeal marker genes from 316 MAGs were generated via GTDB-Tk’s identify and align workflow. IQ-TREE v2.2.0^54^ was used to construct a maximum-likelihood tree. The resultant tree was visualized with the Interactive Tree of Life (iTOL) v5.0 online tool^55^.

### Functional annotations

To characterize the functional potential of all MAGs from deep-sea sediment samples, we performed functional annotation using METABOLIC v4.0^56^. To obtain representative coding sequences, predicted CDSs were pooled and clustered at 95% nucleotide similarity with CD-HIT v4.8.1^57^. Gene abundance was quantified by mapping metagenomic reads to this catalog using Salmon v1.5.0^58^. MAGs were annotated using DRAM^59^ with default parameters against KOfam, MEROPS, and dbCAN databases, identifying peptidases, CAZymes, nucleases, lipases, and other proteins.

### Metatranscriptomic analysis

To determine the *in situ* gene expression profiles of CAZymes within MAGs from sediment samples (2022 Site F cold seep 1, 2022 Site F cold seep 2, 2022 Haima cold seep, and 2022 Zhenbei seamount), we conducted metatranscriptomic analysis. Three independent samples were selected for each site. For metatranscriptomic sequencing analysis, total RNA was extracted using TRIzol® Reagent and its quality was assessed (Agilent 2100 Bioanalyser) and quantified (NanoDrop ND-2000, USA). The comprehensive sequencing and analysis protocols were previously established^52^. Transcripts per million (TPM) was used to indicate the transcript levels for genes encoding CAZymes.

### Enrichment and cultivation of deep-sea PVC bacteria

To cultivate deep-sea PVC bacteria, sediment samples were added to 10 mL basal medium supplemented with 3.0 g/L laminarin. The basal medium consisted of 1.0 g/L NH_4_Cl, 1.0 g/L CH_3_COONa, 1.0 g/L NaHCO_3_, 0.5 g/L MgSO_4_, 500 µL/L of 0.1% (w/v) resazurin, 0.7 g/L cysteine, 1.0 L seawater, and adjusted to pH 7.0. These enrichment cultures were incubated anaerobically at 28°C for one month in N_2_-gassed, autoclaved medium, followed by single-colony isolation on agar-spread Hungate tubes. Pure cultures were established and verified by transmission electron microscopy (TEM) and 16S rRNA gene sequencing^10^.

### TEM observation

For morphological observation of strains WC338 and WC36, cell pellets were obtained by centrifuging cultures grown in basal medium with 3.0 g/L laminarin (5,000 × *g*, 10 min). Cell suspensions were adsorbed onto copper grids for 20 min, washed with 10 mM PBS (pH 7.4), and air-dried. Transmission electron micrographs were acquired using a Hitachi HT7700 with the JEOL JEM 12000 EX at 100 kV.

### Phylogenetic analysis

Full-length 16S rRNA gene sequences from strains WC338, WC36, WC47, WC29 (obtained from their genomes), and related taxa (from NCBI) were aligned using MAFFT v7.0^60^ and manually curated. Phylogenetic trees were constructed using the maximum likelihood method via the W-IQ-TREE web server (http://iqtree.cibiv.univie.ac.at)^61^. The final tree was edited and visualized with the Interactive Tree of Life (iTOL) v5.0 online tool^55^.

### Growth assay

To assess laminarin’s effect on strains WC338 and WC36, 100 µL of fresh cultures were inoculated into 10 mL basal medium with or without 3.0 g/L laminarin in triplicate Hungate tubes. Cultures were incubated anaerobically at 28°C. Growth was monitored daily by measuring OD_600_ using a microplate reader (Tecan Infinite M1000 Pro, Switzerland) until the stationary phase was reached.

### Determination of laminarin content

Total laminarin content was determined using a modified phenol-sulfuric acid colorimetric assay^62^. Briefly, a standard curve was generated using known concentrations of glucose (0-100 µg/mL) as the standard. Subsequently, aliquots (2 mL) of culture supernatant were collected daily from the initial inoculation up to day 9 of incubation. Each experimental condition was performed with three biological replicates. For each sample (both glucose standards and culture supernatants), 2 mL sample was combined with 1 mL of 5% phenol, and 5 mL of concentrated sulfuric acid was rapidly introduced. The mixture was vortexed for 5 s, incubated at 37 °C for 30 min, and its absorbance measured at 490 nm after cooling to room temperature. The laminarin content was calculated based on the absorbance values and the established standard curve.

### Transcriptomic analysis

Strains WC338 and WC36 were cultured in basal medium supplemented with either 0.05 g/L or 3.0 g/L laminarin for 5 days to evaluate transcriptional effects. Cell suspensions were harvested from these cultures for subsequent transcriptomic analysis. Detailed methods for transcriptomic sequencing analysis are described previously^10^.

## Data availability

Raw amplicon sequencing data for seven deep-sea sediment samples are available under NCBI SRA accession numbers PRJNA688815 and PRJNA1281722 (previously described). The 316 deep-sea MAGs presented in this study are accessible via BioProject accession numbers PRJNA1337902 and PRJNA1337911. Complete genome sequences for strains WC338 (CP194516) and WC36 (CP085689), and their respective 16S rRNA gene sequences (WC338: PV668772; WC36: OK614042; WC47: PV682642; WC29: PV682643), are deposited in GenBank. Raw transcriptomic reads for strains WC338 and WC36 are available under NCBI SRA accession numbers PRJNA1337914 and PRJNA1337918.

## Acknowledgements

This work was supported by the National Natural Science Foundation of China (Grant Nos. 42406104 and 42530409), the NSFC Innovative Group Grant (No. 42221005), Science and Technology Innovation Project of Laoshan Laboratory (Grant Nos. LSKJ202203103 and 2022QNLM030004-3), Shandong Provincial Natural Science Foundation (Grant Nos. ZR2024ZD49 and ZR2023QD010), Major Research Plan of the National Natural Science Foundation (Grant No. 92351301), Qingdao West Coast New District University Presidents Fund (Grant No. E42424101N), and Taishan Scholars Program (Grant Nos. tstp20230637 and tsqn202312264). We extend special appreciation to the captain and crew of the *RV KEXUE*, as well as the FAXIAN ROV team for assistance with sample collection.

## Author contributions

RZ and CS conceived and designed the study; RZ and CW conducted most of the experiments; RZ and CW collaboratively analyzed the omics data; RZ, CW, and CS lead the writing of the manuscript; all authors contributed to and reviewed the manuscript.

## Competing interests

The authors declare that they have no any competing interests.

---

**Extended Data Figure 1.**
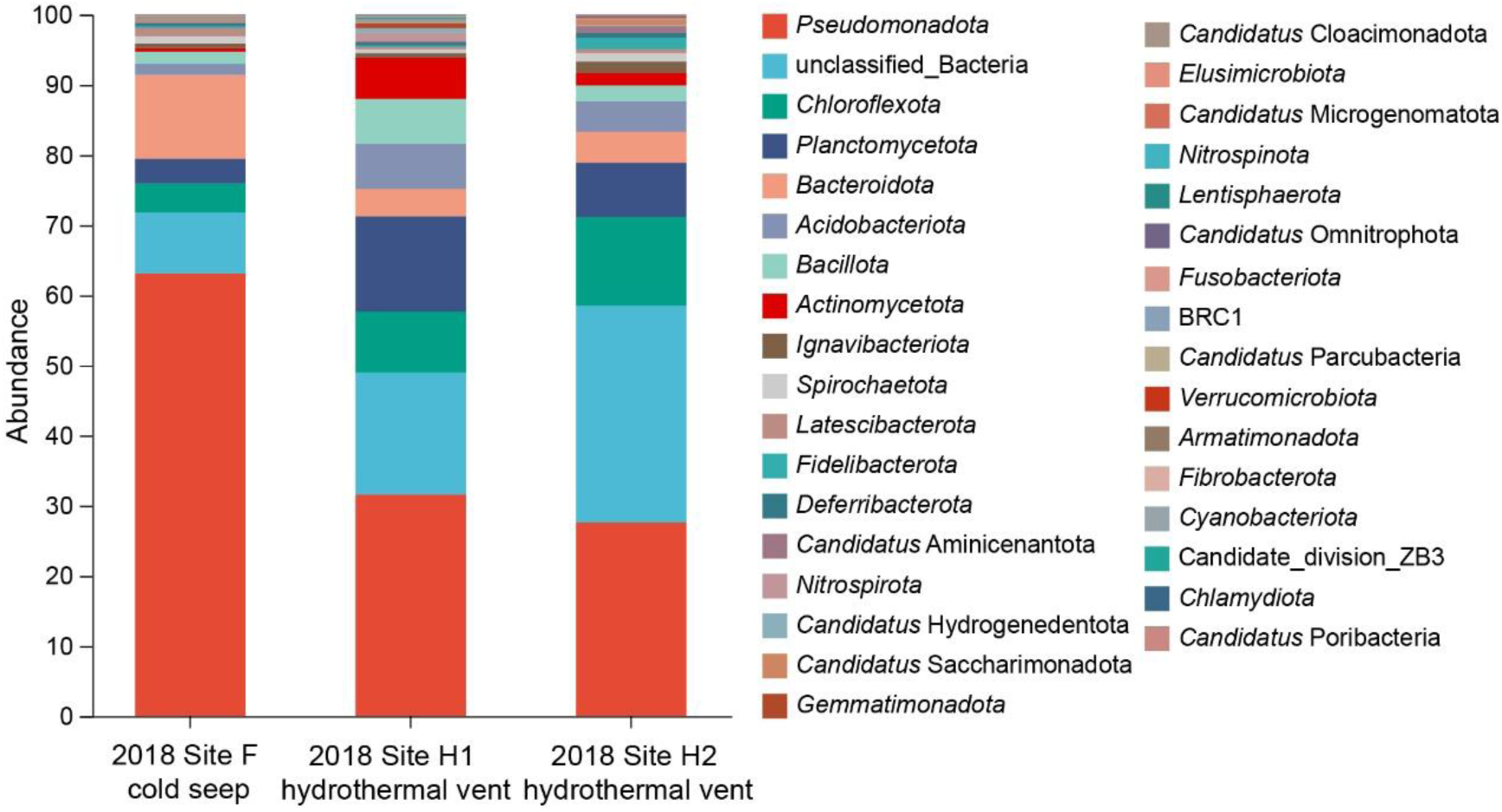
Relative abundances of dominant prokaryotic communities in deep-sea cold seep and hydrothermal vent sediments (collected in 2018) at the phylum level.

**Extended Data Figure 2.**
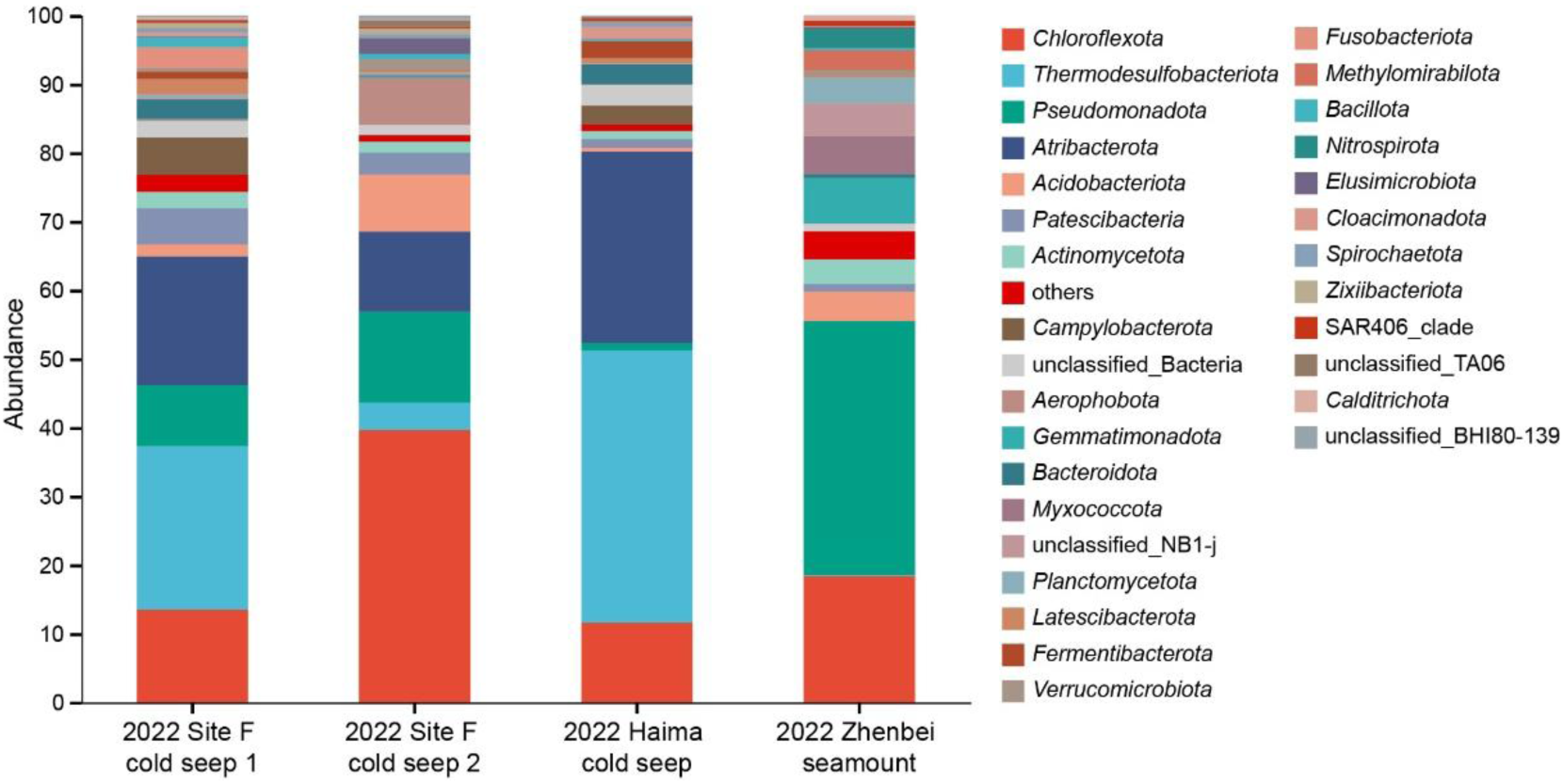
Relative abundances of dominant prokaryotic communities in deep-sea cold seep and seamount sediments (collected in 2022) at the phylum level.

**Extended Data Figure 3.**
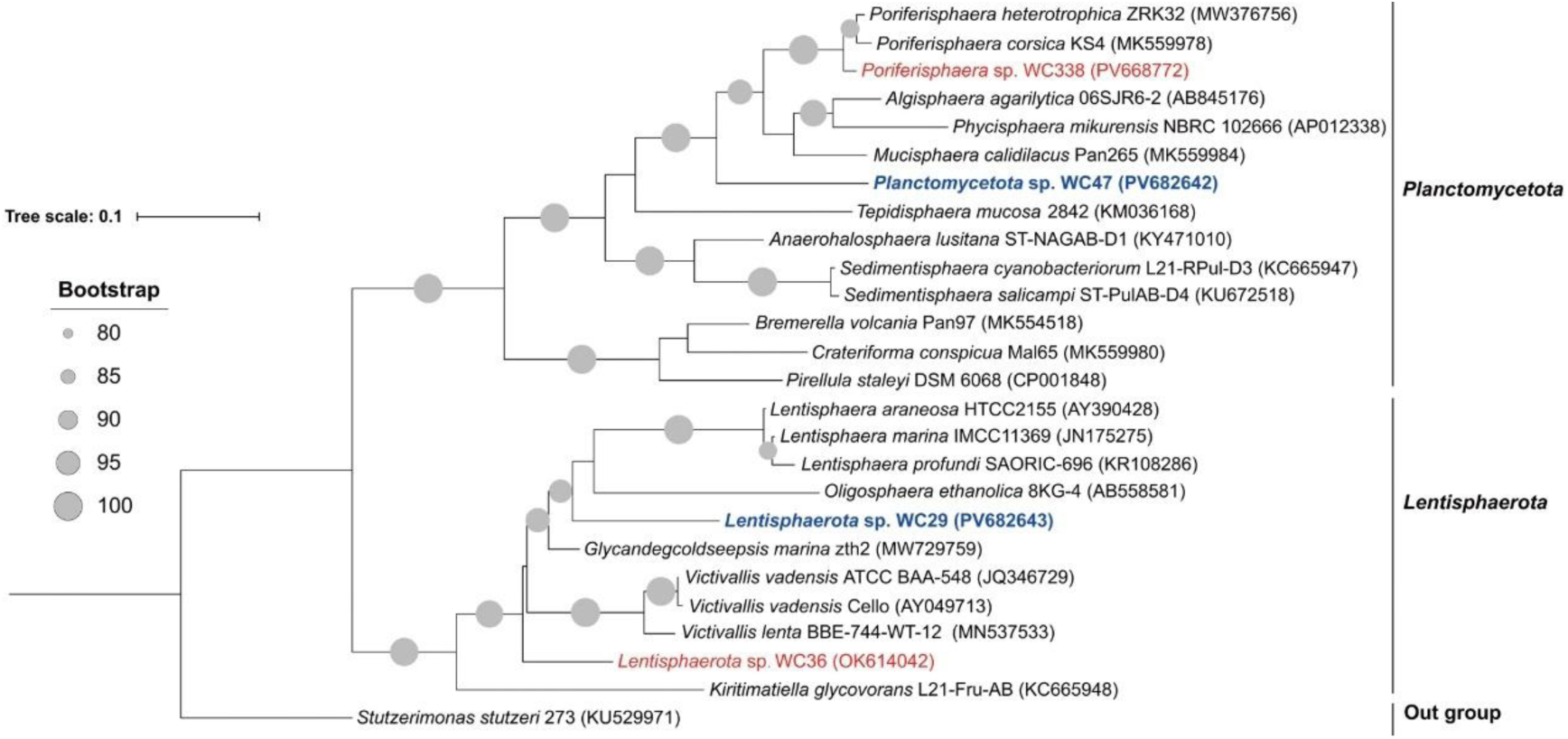
Laminarin facilitates the isolation and cultivation of PVC superphylum bacteria from deep-sea sediments. Maximum-likelihood phylogenetic tree of cultured PVC superphylum isolates include *Planctomycetota* strain WC47 (hydrothermal vent) and *Lentisphaerota* strain WC29 (seamount), based on 16S rRNA gene sequences. Bar, 0.1 substitutions per nucleotide position.

## Notes

### Competing Interest Statement

The authors have declared no competing interest.

## References

1 Jørgensen, B. B. & Boetius, A. Feast and famine--microbial life in the deep-sea bed. Nat. Rev. Microbiol. 5, 770–781, (2007).

2 Horsfield, B. et al. Living microbial ecosystems within the active zone of catagenesis: Implications for feeding the deep biosphere. Earth Planet Sci. Lett. 246, 55–69, (2006).

3 Dong, X. et al. Metabolic potential of uncultured bacteria and archaea associated with petroleum seepage in deep-sea sediments. Nat. Commun. 10, 1816, (2019).

4 Dombrowski, N., Teske, A. P. & Baker, B. J. Expansive microbial metabolic versatility and biodiversity in dynamic Guaymas Basin hydrothermal sediments. Nat. Commun. 9, 4999, (2018).

5 Pérez Castro, S., et al. Degradation of biological macromolecules supports uncultured microbial populations in Guaymas Basin hydrothermal sediments. ISME J. 15, 3480–3497, (2021).

6 Jørgensen, B. B., Andrén, T. & Marshall, I. P. G. Sub-seafloor biogeochemical processes and microbial life in the Baltic Sea. Environ. Microbiol. 22, 1688–1706, (2020).

7 Orsi, W. D., Richards, T. A. & Francis, W. R. Predicted microbial secretomes and their target substrates in marine sediment. Nat. Microbiol. 3, 32–37, (2018).

8 Zheng, R., Cai, R., Liu, R., Liu, G. & Sun, C. *Maribellus comscasis* sp. nov., a novel deep-sea *Bacteroidetes* bacterium, possessing a prominent capability of degrading cellulose. Environ. Microbiol. 23, 4561–4575, (2021).

9 Zhu, X. Y. et al. Deep-sea *Bacteroidetes* from the Mariana Trench specialize in hemicellulose and pectin degradation typically associated with terrestrial systems. Microbiome 11, 175, (2023).

10 Zheng, R. et al. Characterization of the first cultured free-living representative of *Candidatus* Izemoplasma uncovers its unique biology. ISME J. 15, 2676–2691, (2021).

11 Becker, S., Scheffel, A., Polz, M. F. & Hehemann, J. H. Accurate quantification of laminarin in marine organic matter with enzymes from marine microbes. Appl. Environ. Microbiol. 83, e03389–03316, (2017).

12 Sidhu, C. et al. Dissolved storage glycans shaped the community composition of abundant bacterioplankton clades during a North Sea spring phytoplankton bloom. Microbiome 11, 77, (2023).

13 Alderkamp, A. C., van Rijssel, M. & Bolhuis, H. Characterization of marine bacteria and the activity of their enzyme systems involved in degradation of the algal storage glucan laminarin. FEMS Microbiol. Ecol. 59, 108–117, (2007).

14 Becker, S. et al. Laminarin is a major molecule in the marine carbon cycle. Proc. Natl. Acad. Sci. U. S. A. 117, 6599–6607, (2020).

15 Teeling, H. et al. Substrate-controlled succession of marine bacterioplankton populations induced by a phytoplankton bloom. Science 336, 608–611, (2012).

16 Keith, S. C. & Arnosti, C. Extracellular enzyme activity in a river-bay-shelf transect: variations in polysaccharide hydrolysis rates with substrate and size class. Aquat Microb. Ecol. 24, 243–253, (2001).

17 Arnosti, C., Durkin, S. & Jeffrey, W. H. Patterns of extracellular enzyme activities among pelagic marine microbial communities: implications for cycling of dissolved organic carbon. Aquat Microb. Ecol. 38, 135–145, (2005).

18 Vigneron, A. et al. Comparative metagenomics of hydrocarbon and methane seeps of the Gulf of Mexico. Sci. Rep. 7, 16015, (2017).

19 Rivas-Marin, E. & Devos, D. P. The Paradigms They Are a-Changin’: past, present and future of PVC bacteria research. Anton Leeuw Int J G 111, 785–799, (2018).

20 Devos, D. P. & Ward, N. L. Mind the PVCs. Environ. Microbiol. 16, 1217–1221, (2014).

21 Devos, D. P. Reconciling Asgardarchaeota phylogenetic proximity to eukaryotes and *Planctomycetes* cellular features in the evolution of life. Mol. Biol. Evol. 38, 3531–3542, (2021).

22 Pollet, T., Tadonléké, R. D. & Humbert, J. F. Spatiotemporal changes in the structure and composition of a less-abundant bacterial phylum (*Planctomycetes*) in two perialpine lakes. Appl. Environ. Microbiol. 77, 4811–4821, (2011).

23 Dedysh, S. N. & Ivanova, A. A. *Planctomycetes* in boreal and subarctic wetlands: diversity patterns and potential ecological functions. FEMS Microbiol. Ecol. 95, (2019).

24 Buckley, D. H., Huangyutitham, V., Nelson, T. A., Rumberger, A. & Thies, J. E. Diversity of *Planctomycetes* in soil in relation to soil history and environmental heterogeneity. Appl. Environ. Microbiol. 72, 4522–4531, (2006).

25 Woebken, D. et al. Fosmids of novel marine *Planctomycetes* from the Namibian and Oregon coast upwelling systems and their cross-comparison with *planctomycete* genomes. ISME J. 1, 419–435, (2007).

26 Goffredi, S. K. & Orphan, V. J. Bacterial community shifts in taxa and diversity in response to localized organic loading in the deep sea. Environ. Microbiol. 12, 344–363, (2010).

27 Wiegand, S., Jogler, M. & Jogler, C. On the maverick *Planctomycetes*. FEMS Microbiol. Rev. 42, 739–760, (2018).

28 Wiegand, S. et al. Cultivation and functional characterization of 79 planctomycetes uncovers their unique biology. Nat. Microbiol. 5, 126–140, (2020).

29 Wang, J., et al. *Gimesia benthica* sp. nov., a planctomycete isolated from a deep-sea water sample of the Northwest Indian Ocean. Int. J. Syst. Evol. Microbiol. 70, 4384–4389, (2020).

30 Zheng, R., Wang, C., Liu, R., Cai, R. & Sun, C. Physiological and metabolic insights into the first cultured anaerobic representative of deep-sea Planctomycetes bacteria. Elife 12, (2024).

31 Kulichevskaya, I. S., et al. *Frigoriglobus tundricola* gen. nov., sp. nov., a psychrotolerant cellulolytic planctomycete of the family Gemmataceae from a littoral tundra wetland. Syst. Appl. Microbiol. 43, 126129, (2020).

32 Achberger, A. M. et al. Inactive hydrothermal vent microbial communities are important contributors to deep ocean primary productivity. Nat. Microbiol. 9, 657–668, (2024).

33 Herndl, G. J. & Reinthaler, T. Microbial control of the dark end of the biological pump. Nat. Geosci. 6, 718–724, (2013).

34 Zheng, R., Cai, R., Wang, C., Liu, R. & Sun, C. Characterization of the first cultured representative of “*Candidatus* Thermofonsia” clade 2 within *Chloroflexi* reveals its phototrophic lifestyle. mBio 13, e0028722, (2022).

35 Dong, X. et al. Evolutionary ecology of microbial populations inhabiting deep sea sediments associated with cold seeps. Nat. Commun. 14, 1127, (2023).

36 Sun, T., Wu, D., Wu, N. & Yin, P. The effects of organic matter and anaerobic oxidation of methane on the microbial sulfate reduction in cold seeps. Front. Mar. Sci. 10, (2023).

37 Bowers, R. M. et al. Corrigendum: Minimum information about a single amplified genome (MISAG) and a metagenome-assembled genome (MIMAG) of bacteria and archaea. Nat. Biotechnol. 36, 660, (2018).

38 Cai, R., Zhang, J., Liu, R. & Sun, C. Metagenomic insights into the metabolic and ecological functions of abundant deep-sea hydrothermal vent DPANN archaea. Appl. Environ. Microbiol. 87, e03009–03020, (2021).

39 Zhang, C. et al. The majority of microorganisms in gas hydrate-bearing subseafloor sediments ferment macromolecules. Microbiome 11, 37, (2023).

40 Oni, O. E. et al. Microbial Communities and Organic Matter Composition in Surface and Subsurface Sediments of the Helgoland Mud Area, North Sea. Front. Microbiol. 6, 1290, (2015).

41 Dell’Anno, A. & Danovaro, R. Extracellular DNA plays a key role in deep-sea ecosystem functioning. Science 309, 2179, (2005).

42 Dang, Y. R. et al. Phytoplankton-derived polysaccharides and microbial peptidoglycans are key nutrients for deep-sea microbes in the Mariana Trench. Microbiome 12, 77, (2024).

43 Klimek, D., Herold, M. & Calusinska, M. Comparative genomic analysis of *Planctomycetota* potential for polysaccharide degradation identifies biotechnologically relevant microbes. BMC Genomics 25, 523, (2024).

44 Pérez-Cruz, C. et al. Mechanisms of recalcitrant fucoidan breakdown in marine Planctomycetota. Nat. Commun. 15, 10906, (2024).

45 Kim, M., Oh, H. S., Park, S. C. & Chun, J. Towards a taxonomic coherence between average nucleotide identity and 16S rRNA gene sequence similarity for species demarcation of prokaryotes. Int. J. Syst. Evol. Microbiol. 64, 346–351, (2014).

46 Yarza, P. et al. Uniting the classification of cultured and uncultured bacteria and archaea using 16S rRNA gene sequences. Nat. Rev. Microbiol. 12, 635–645, (2014).

47 Clerc, E. E. et al. Strong chemotaxis by marine bacteria towards polysaccharides is enhanced by the abundant organosulfur compound DMSP. Nat. Commun. 14, 8080, (2023).

48 Li, D. et al. Biochemical insights into a novel family 2 glycoside hydrolase with both β-1,3-galactosidase and β-1,4-galactosidase activity from the Arctic. Mar. Drugs 21, 521, (2023).

49 Liberato, M. V. et al. Insights into the dual cleavage activity of the GH16 laminarinase enzyme class on β-1,3 and β-1,4 glycosidic bonds. J. Biol. Chem. 296, 100385, (2021).

50 Choi, A., Song, J., Joung, Y., Kogure, K. & Cho, J. C. *Lentisphaera profundi* sp. nov., isolated from deep-sea water. Int. J. Syst. Evol. Microbiol. 65, 4186–4190, (2015).

51 Xiong, J. et al. Effects of menopausal hormone therapy on gut microbiota in postmenopausal women and the relationship with bone metabolism. Front. Med. 12, 1682925, (2025).

52. Wang, C., Zheng, R. & Sun, C. Characterization of deep-sea viruses reveals their unexpected diversity and role in facilitating host metabolism of complex organic matter. bioRxiv, 2025.2003.2021.644689, (2025).

53 Chaumeil, P. A., Mussig, A. J., Hugenholtz, P. & Parks, D. H. GTDB-Tk: a toolkit to classify genomes with the Genome Taxonomy Database. Bioinformatics 36, 1925–1927, (2019).

54 Minh, B. Q. et al. IQ-TREE 2: new models and efficient methods for phylogenetic inference in the genomic Era. Mol. Biol. Evol. 37, 1530–1534, (2020).

55 Letunic, I. & Bork, P. Interactive Tree Of Life (iTOL) v5: an online tool for phylogenetic tree display and annotation. Nucleic Acids Res. 49, W293–W296, (2021).

56 Zhou, Z. et al. METABOLIC: high-throughput profiling of microbial genomes for functional traits, metabolism, biogeochemistry, and community-scale functional networks. Microbiome 10, 33, (2022).

57 Fu, L., Niu, B., Zhu, Z., Wu, S. & Li, W. CD-HIT: accelerated for clustering the next-generation sequencing data. Bioinformatics 28, 3150–3152, (2012).

58 Patro, R., Duggal, G., Love, M. I., Irizarry, R. A. & Kingsford, C. Salmon provides fast and bias-aware quantification of transcript expression. Nat. Methods 14, 417–419, (2017).

59 Shaffer, M. et al. DRAM for distilling microbial metabolism to automate the curation of microbiome function. Nucleic Acids Res. 48, 8883–8900, (2020).

60 Katoh, K., Rozewicki, J. & Yamada, K. D. MAFFT online service: multiple sequence alignment, interactive sequence choice and visualization. Brief Bioinform. 20, 1160–1166, (2019).

61 Trifinopoulos, J., Nguyen, L. T., von Haeseler, A. & Minh, B. Q. W-IQ-TREE: a fast online phylogenetic tool for maximum likelihood analysis. Nucleic Acids Res. 44, W232–W235, (2016).

62 Yue, F. et al. Effects of monosaccharide composition on quantitative analysis of total sugar content by phenol-sulfuric acid method. Front. Nutr. 9, 963318, (2022).

